# Therapeutic antibodies, targeting the SARS-CoV-2 spike N-terminal domain, protect lethally infected K18-hACE2 mice

**DOI:** 10.1101/2021.02.02.428995

**Authors:** Tal Noy-Porat, Adva Mechaly, Yinon Levy, Efi Makdasi, Ron Alcalay, David Gur, Moshe Aftalion, Reut Falach, Shani Leviatan Ben-Arye, Shirley Lazar, Ayelet Zauberman, Eyal Epstein, Theodor Chitlaru, Shay Weiss, Hagit Achdout, Jonathan D. Edgeworth, Raghavendra Kikkeri, Hai Yu, Xi Chen, Shmuel Yitzhaki, Shmuel C. Shapira, Vered Padler-Karavani, Ohad Mazor, Ronit Rosenfeld

## Abstract

Since the onset of the current COVID-19 pandemic, high priority is given to the development of neutralizing antibodies, as a key approach for the design of therapeutic strategies to countermeasure and eradicate the disease. Previously, we reported the development of human therapeutic monoclonal antibodies (mAbs) exhibiting very high protective ability. These mAbs recognize epitopes on the spike receptor binding domain (RBD) of SARS-CoV-2 that is considered to represent the main rout of receptor engagement by the SARS-CoV-2 virus. The recent emergence of viral variants emphasizes the notion that efficient antibody treatments need to rely on mAbs against several distinct key epitopes in order to circumvent the occurrence of therapy escape-mutants. Here we report the isolation and characterization of 12 neutralizing mAbs, identified by screening a phage-display library constructed from lymphatic cells collected from severe COVID-19 patients. The antibodies target three distinct epitopes on the spike N-terminal domain (NTD) of SARS-CoV-2, one of them defining a major site of vulnerability of the virus. Extensive characterization of these mAbs suggests a neutralization mechanism which relies both on amino-acid and *N*-glycan recognition on the virus, and involvement of receptors other than the hACE2 on the target cell. Two of the selected mAbs, which demonstrated superior neutralization potency *in vitro*, were further evaluated *in vivo*, demonstrating their ability to fully protect K18-hACE2 transgenic mice even when administered at low doses and late after infection. The study demonstrates the high potential of the mAbs for therapy of SARS-CoV-2 infection and underlines the possible role of the NTD in mediating infection of host cells via alternative cellular portals other than the canonical ACE2 receptor.

## Introduction

The ongoing COVID-19 pandemic caused by the severe acute respiratory syndrome coronavirus 2 (SARS-CoV-2) inflicted an enormous public health and economic crisis and continues to spread globally. Despite the recent onset of mass vaccination campaigns, the pandemic exhibits unprecedented extent of morbidity and mortality. Consequently, there is still an urgent need for effective therapeutics. The virus spike (S) glycoprotein, essential for the recognition of its human cognate angiotensin-converting enzyme 2 (hACE2) receptor on target cells and subsequent viral internalization, was implicated as the main target for coronavirus neutralizing monoclonal antibodies (mAbs). Processing of the S protein by host proteases results in the generation of the S1 subunit, responsible for receptor engagement and the S2 subunit which mediates membrane fusion^1^. Two functional domains can be distinguished within the S1 subunit: the N-terminal domain (NTD) and the C-terminal domain (CTD)^2^. SARS-CoV-2, similar to SARS-CoV, uses CTD as the receptor binding domain (RBD), associated with direct contact to the hACE2 receptor^3,4^. However, some other betacoronaviruses use the NTD of their S1 subunit to bind various host cell receptors. For example, the HCoV-OC43 and HCoV-HKU1 respective NTDs recognize terminal 9-*O*-acetylated sialic acid moieties of glycosylated cell-surface receptors^5–7^. Similarly, the betacoronavirus murine hepatitis virus uses its NTD for binding the protein receptor CEACAM1^8^ and MERS-CoV utilizes a dual-receptor strategy, using both protein (DPP4) and sialoside receptors to infect host cells^5,9^. The role of the NTD of SARS-CoV-2 is yet to be fully understood, but accumulating evidence suggests that it is important for SARS-CoV-2 infectivity. Comparison of the NTD primary sequences and spatial architectures of SARS-CoV, MERS-CoV and SARS-CoV-2, reveals several loop regions common to both SARS-CoV-2 and MERS-CoV but lacking in SARS-CoV^10^. These loops constitute a potential sialoside binding pocket and therefore may use human sialosides as an alternate receptors for SARS-CoV-2 infection in a variety of tissues. Moreover, L-SIGN and DC-SIGN, members of the C-type lectin superfamily, were recently suggested as possible alternative receptors for SARS-CoV-2, via interaction with the NTD domain^11–13^. Theoretically, the wide repertoire of human tissues and immune cells expressing these receptors, may explain the presence of SARS-CoV-2 in many tissues of the infected host which do not exhibit significant or even detectable ACE2 expression levels.

Direct interference of the SARS-CoV-2 engagement with its ACE2 cognate receptor on target cells, represents the main mechanism of virus neutralization, exhibited by most of the reported therapeutic antibodies developed for COVID-19 treatment. Accordingly, these antibodies were identified on the basis of their ability to bind RBD, some of which present substantial neutralization efficiencies^14–19^. Nevertheless, one of the major concerns when using therapeutic anti-viral antibodies is the emergence of escape-variants, due to the rapid mutation frequency of viral pathogens and the selective pressure imposed upon one defined domain by a single mAb, recognizing one particular epitope. One of the most effective ways to circumvent this therapeutic complication is treatment with mAbs directed to different epitopes. Indeed, administration of mAbs combination was shown to efficiently reduce the development of antibody resistance in the case of HIV infection^20^ and also in the case of SARS-CoV-2^21^. Therefore, development of neutralizing mAbs targeting diverse epitopes on the S glycoprotein represents a high priority objective.

So far, only few reports documented isolation of NTD-specific human mAbs^15,22–24^. The mechanism of neutralization promoted by anti-NTD mAbs is not fully understood and in general, these reported antibodies exhibited inferior neutralizing capabilities compared to RBD-specific mAbs. However, it seems that the spike NTD represents also an important element involved in SARS-CoV-2 infectivity, and therefore, a possible additional target for neutralizing antibodies. The recent emergence of escape variants further encouraged consideration of neutralizing mAbs against NTD as a therapeutic strategy, possibly in combination with anti-RBD mAbs.

Accordingly, the objective of the study reported here was isolation of potent neutralizing anti-NTD mAbs, by a phage-display approach, using libraries constructed from lymphocytes collected from human severe COVID-19 patients.

## Methods

### Recombinant proteins

The SARS-CoV-2 spike (S) stabilized soluble ectodomain, S1 subunit, N-terminal domain (NTD), receptor binding domain (RBD) and the human angiotensin-converting enzyme 2 (hACE2) were produced as previously described^19^. L-SIGN-hFc tag was purchased from R&D systems (USA). Antibody 4A8 was produced in-house according to GenBank accession no. MT622682/3.

### Blood samples

Anonymized surplus blood samples of COVID19 patients with persistent critical illness >21 days after first day of symptom onset and taken as part of routine care (St Thomas’ Hospital, London, UK) were retrieved at the point of being discarded. Work was undertaken in accordance with UK Research Ethics Committee (REC) approval for laboratory research projects investigating the immune response to COVID-19 infection with academic collaborators (REC reference 20/SC/0310). Samples were treated in accordance with the biosafety guidelines of the IIBR in BL3 facility. PBMCs were separated from fresh whole blood using density centrifugation by Ficoll. Sera samples were heat-inactivated (20 min at 60°C) prior to use for binding or neutralizing assays.

### Phage display scFv libraries construction and antibodies selection

Total RNA was purified from PBMCs using RNeasy mini kit (Qiagen GmbH, Germany) and used for cDNA synthesis (Verso cDNA synthesis kit; Thermoscientific, USA). Antibodies variable region coding fragments were amplified directly from cDNA, using a human V/J specific set of primers^19^. Phage-display library was constructed as previously detailed^19^. Briefly, heavy and light Ig variable domains (V_H_ and V_L_) were amplified and used in PCR overlap extension reaction, to obtain scFv repertoire cloned into pCC16 phagemid vector^56^, using *Nco*I/*Not*I. Total of 1.3×10^9^ independent clones obtained, representing the library complexity.

For phage production, 25 ml of logarithmic bacteria culture (OD_600_=0.6) in YPD supplemented with 100 µg/ml ampicillin and 1% glucose (YPD-Amp-Glu) were infected with M13KO7 helper phage (New England Biolabs, USA) at 7×10^9^ plaque forming unit (PFU) per ml (~1:20 multiplicity of infection) by incubating at 37 °C for 30 min without shaking, followed by 30 min at 120 rpm. Infected cells were harvested by centrifugation (1800 g for 5 min) and resuspended in 100 ml YPD supplemented with 100 µg/ml ampicillin and 50 µg/ml kanamycin. After overnight growth at 30 °C at 200 rpm, the cells were removed by centrifugation (1800 g at 4 °C for 10 min). The culture supernatant containing the phages was filtered through a 0.45-µm filter and then precipitated with 1/5 volume of 20% PEG6000 (polyethylene glycol) in a 2.5 M NaCl solution, for 2 h on ice. Phage particles were pelleted by centrifugation (9000 *g* at 4 °C for 1 h) and re-dissolved in 5 ml Dulbecco’s Phosphate Buffered Saline (PBS; Biological Industries, Israel).

Panning was performed against monomeric NTD directly absorbed to polystyrene plates (Maxisorb 96-well microtiter plates; Nunc, Sigma-Aldrich, USA). Blocking of plates and phages was conducted for 60 min using two blocking solutions: 3% BSA (in PBS) or 2% skimmed milk and 0.05% Tween20 in PBS. The blocking solutions were alternated between panning cycles. All washing steps were performed using PBST (PBS containing 0.05% Tween20 and 2% BSA). For each panning cycle, 5 µg/ml antigen was used to coat the polystyrene plate and after an overnight incubation, plates were washed and blocked. For the first panning cycle, approximately 1×10^11^ phages were incubated with the antigen coated plates for 60 min, followed by a total of 3 washes with PBST. Phages were eluted by incubation with 1ml of 100 mM Triethylamine (Sigma, Israel) for 30 min and following neutralization (in 200 µl 1M Tris-HCl, pH=7.4), were used to infect 5 ml of *E. coli* TG1 strain, by incubation at 37°C for 30 min without shaking followed by 30 min at 120 rpm. The bacterial culture was plated on YPD-Amp-Glu agar, and incubated overnight at 30°C. Clones were harvested into 5 ml YPD-Amp-Glu with 20% glycerol solution and phage production for the next round of panning was conducted in 10 ml medium, as described above. Two additional panning cycles were conducted essentially similarly, with the following modifications: 10^10^ phages were used as input for the 2^nd^ and 3^rd^ cycles, and the washing steps were increased to include 6 washes of PBST.

Single colonies were randomly picked from the third cycle output and screen of specific binders was performed, using phage ELISA against NTD Vs RBD.

### Single chain Fv (scFv) individual clone diversity and sequence verification

TAB-RI_For (CCATGATTACGCCAAGCTTTGGAGCC) and CBD-AS_Rev (GAATTC AACCTTCAAATTGCC) phagemid specific primers were used for colony PCR and sequence analysis of scFv antibody individual clones. Colony PCR products, were analyzed on 1.5 % agarose gel, to confirm the intact of the scFv. Restriction fragment size polymorphism (RFLP), was performed using *MvaI* (FastDigest; Thermoscientific, USA) to evaluate sequence variability of scFv individual clones. Following colony PCR, 5µl of the PCR products were taken directly for restriction with 0.5 µl *Mva*I and 1 µl buffer x10 (provided by the manufacturer) in a 10 µl reaction volume. Restriction was conducted for one h at 37°C, and the entire reaction mix was then resolved on 3% agarose gel. Nucleic acid sequence analysis of individual scFv fragments, was performed to the colony PCR product, using SeqStudio Genetic Analyzer (Applied Biosystems, USA).

### Production of Antibodies

Phagemid DNA of the desired clones were isolated using QIAprep spin Miniprep kit (Qiagen, GmbH, Hilden, Germany), and the entire scFv was cloned into a pcDNA3.1+ based expression vector that was modified, providing the scFv with the human (IgG1) CH2-CH3 Fc fragments, resulting in scFv-Fc antibody format. For IgG-YTE format, VH and VL fragments of each antibody were separately amplified and cloned into a pcDNA3.1+ based expression vector including either full IgG1 constant region (carrying YTE mutations^14^) for the heavy variable fragments, or Kappa/Lambda constant regions for the light variable fragments. All antibody formats were expressed using ExpiCHOTM Expression system (Thermoscientific, USA) and purified on HiTrap Protein-A column (GE healthcare, UK). The integrity and purity of the antibodies were analyzed using SDS-PAGE.

### ELISA

Direct ELISA^57^ consisted of coating microtiter plates with 1 μg/ml of recombinant SARS-CoV-2 spike, S1 domain, RBD or NTD subunits. For phage ELISA, HRP-conjugated anti-M13 antibody (Sino Biological, USA, Cat# 11973-MM05T-H lot HO13AU601; used at 1:8000 working dilution) was used following detection with TMB substrate (Millipore, USA). ELISA of both sera and recombinant human antibodies was applied with AP-conjugated Donkey anti-human IgG (Jackson ImmunoResearch, USA, Cat# 709-055-149 lot 130049; used at 1:2000 working dilution) following detection using *p*-nitrophenyl phosphate (*p*NPP) substrate (Sigma, Israel).

In order to evaluate sialoglycans involvement in the anti-NTD antibodies binding, Costar 96-well were coated overnight at 4°C with 0.1 µg/well NTD or RBD in coating buffer (50 mM sodium carbonate-bicarbonate buffer, pH 9.5). Wells were blocked for 2 hours at RT with blocking solution (PBS with 1% chicken ovalbumin; Sigma). For ELISA inhibition assay, primary antibodies (1.5 µg/ml in blocking solution) were pre-incubated for two hours on ice in the presence or absence of inhibitors [blocking buffer for no-inhibition control, 8 mM 2-*O*-methyl-α-Neu5Ac (Ac2Me), 8 mM D-Glucuronic acid (GlcA; Sigma-Aldrich), or with 0.06 mM sialoglycopeptides (GP) produced from Neu5Gc-deficient *Cmah*^−*/*−^ mice sera^33^ containing 3.4 mM Neu5Ac-GP (validated by DMB-HPLC)]. This mAb-inhibitor pre-incubated mixture was added to the coated plate and incubated at RT for 2 hours. Wells were washed three times with PBST, following 1 hour incubation at RT with HRP-goat-anti-human IgG H+L (Jackson ImmunoResearch, USA, Cat#109-035-003 lot 149724; used at final concentration of 0.1µg/ml). After washing three times with PBST (PBS with 0.1% Tween-20), wells were developed with 0.5 mg/ml *O*-phenylenediamine in citrate-PO_4_ buffer, pH 5.5, reaction was stopped with H_2_SO_4_ and absorbance was measured at 490 nm wavelength on SpectraMax M3 (Molecular Devices).

The endogenous humoral response was tested in mice sera samples collected at 21 days post infection (dpi), evaluated by ELISA against SARS-CoV-2 spike, NTD and RBD essentially as described above, using AP-conjugated Donkey anti-mouse IgG (H+L) minimal cross (Jackson ImmunoResearch, USA, Cat# 715-055-150, lot 142717) used at 1:2,000.

### Biolayer interferometry for affinity measurements and epitope binning

Binding studies were carried out using the Octet system (ForteBio, USA, Version 8.1, 2015) that measures biolayer interferometry (BLI). All steps were performed at 30°C with shaking at 1500 rpm in a black 96-well plate containing 200 μl solution in each well. Streptavidin-coated biosensors were loaded with biotinylated antibody (5-10 µg/ml) to reach 1 nm wavelength shift followed by a wash. The sensors were then reacted for 300 s with increasing concentrations of monomeric NTD (association phase) and then transferred to buffer-containing wells for another 600 s (dissociation phase). Binding and dissociation were measured as changes over time in light interference after subtraction of parallel measurements from unloaded biosensors. Sensorgrams were fitted with a 1:1 binding model (Supplementary Fig. 1) using the Octet data analysis software 8.1 (Fortebio, USA, 2015).

For the binning experiments of antibodies pairs, antibody-loaded sensors were incubated with a fixed concentration of S1 (20 µg/ml), washed and incubated with the non-labeled antibody counterpart. In each set of experiments, the background signal was obtained from a parallel sensor incubated with the homologous antibody and sensograms are presented after subtraction of the background signal (Supplementary Fig. 2).

To study the binding of hACE2 to S1 in the presence of antibodies, antibody-loaded sensors were incubated with S1, washed and incubated with hACE2 (10μg/ml).

In order to study the interaction of L-SIGN with NTD, L-SIGN was immobilized on Protein A sensors and allowed to bind NTD or RBD in binding buffer (50mM Hepes, 5 mM CaCl_2_, 5 mM MgCl_2_).

For L-SIGN binding inhibition assay, NTD-His (10 µg/ml) was immobilized on Ni-NTA sensors, incubated with anti-NTD antibodies washed and incubated with L-SIGN (20 µg/ml). No Ab signal was obtained from sensor incubated with L-SIGN without the presence of antibodies and background signal was obtained from a parallel sensor incubated with L-SIGN without the presence of NTD and sensograms are presented after subtraction of the background signal.

### Epitope mapping

Lyophilized 240 biotinylated 15 amino-acid long peptides (with 10 amino-acid overlap) covering the entire ectodomain of SARS-CoV2 spike protein were purchased from JPT (Germany). All peptides were re-suspended in di-methyl sulfoxide (DMSO) to a concentration of 1 mg/ml, aliquoted and stored at −20°C. An aliquot of the peptides was thawed, diluted 1:100 in 1xPBS (to reach 10 µg/ml) and added to Maxisorp ELISA plates (Thermoscientific) pre-coated with streptavidin and blocked with 2% Bovine serum albumin (BSA). Plated peptides were incubated with the individual monoclonal antibodies (5 µg/ml diluted in blocking buffer) and further incubated with donkey anti-human alkaline phosphatase-conjugated secondary antibody (Jackson ImmunoResearch, USA). Immune complexes were identified following incubation with SIGMAFAST™ PNPP (SIGMA) and measuring absorbance at 405 nm. For the modeling of mAbs recognition sites on SARS-CoV-2 S protein, spike structure with PDB ID 7C2L^24^ was used and analyzed by The PyMOL Molecular Graphics System (Version 1.7 Schrödinger, LLC).

### Glycan binding analyses

#### Glycan microarray fabrication

Arrays were fabricated as described^58^ with NanoPrint LM-60 Microarray Printer (Arrayit) on epoxide-derivatized slides (PolyAn 2D) with four 946MP3 pins (5 µm tip, 0.25 µl sample channel, ~100 µm spot diameter; Arrayit) at 16 sub-array blocks on each slide (array V14). Glycoconjugates were distributed into 384-well source plates using 4 replicate wells per sample and 7 µl per well. Each glycoconjugate was printed at 100 µM in an optimized printing buffer (300 mM phosphate buffer, pH 8.4), at four replicate spots. To monitor printing quality, replicate-wells of human IgG (40, 20, 10, 5, and 2.5 ng/µl in PBS with 10% glycerol) and AlexaFlour-555-Hydraside (Invitrogen, at 1 ng/µl in 178 mM phosphate buffer, pH 5.5) were used for each printing run. The humidity level in the arraying chamber was maintained at about 70% during printing. Printed slides were left on an arrayer deck overnight, allowing humidity to drop to ambient levels (40-45%). Next, slides were packed, vacuum-sealed, and stored at room temperature (RT) until used.

#### Sialoglycan microarray binding assay

Slides were developed and analyzed as described^58^ with some modifications. Slides were rehydrated with dH_2_O and incubated for 30 min in a staining dish with 50 °C pre-warmed ethanolamine (0.05 M) in Tris-HCl (0.1 M, pH 9.0) to block the remaining reactive epoxy groups on the slide surface and then washed with 50 °C pre-warmed dH_2_O. Slides were centrifuged at 200 × g for three min and then fitted with ProPlate Multi-Array 16-well slide module (Invitrogen) to divide into the subarrays (blocks). Slides were washed with PBST (0.1% Tween 20), aspirated, and blocked with 200 µl/subarray of blocking buffer (PBS/OVA, 1% w/v ovalbumin, in PBS, pH 7.3) for 1 h at RT with gentle shaking. Next, the blocking solution was aspirated, and 100 µl/block of anti-NTD antibody (5 μg/ml in PBS/OVA) were incubated with gentle shaking for 2 h at RT. Slides were washed four times with PBST and bound antibodies were detected by incubating with secondary antibody (Cy3-goat-anti-human IgG H+L at 1.2 μg/ml; Jackson ImmunoResearch) diluted in PBS/OVA, 200 μl/block at RT for 1 h. Slides were washed three times with PBST and then with PBS for 10 min followed by removal from a ProPlate MultiArray slide module, immediately dipped in a staining dish with dH_2_O for 10 min with shaking, and then centrifuged at 200 × g for 3 min. Dry slides were immediately scanned.

Processed slides were scanned and analyzed as described^58^ at 10 μm resolution with a GenePix 4000B microarray scanner (Molecular Devices) using 350 gain. Image analysis was carried out with Genepix Pro 7.0 analysis software (Molecular Devices). Spots were defined as circular features with a variable radius as determined by the Genepix scanning software. Local background subtraction was performed.

### Cells and virus strains

Vero E6 (ATCC® CRL-1586™) were obtained from the American Type Culture Collection. Cells were grown in Dulbecco’s modified Eagle’s medium (DMEM) supplemented with 10% fetal bovine serum (FBS), MEM non-essential amino acids (NEAA), 2 mM L-glutamine, 100 Units/ml penicillin, 0.1 mg/ml streptomycin and 12.5 Units/ml Nystatin (P/S/N) (Biological Industries, Israel). Cells were cultured at 37°C, 5% CO_2_ at 95% air atmosphere.

ExpiCHO-S (Thermoscientific, USA, Cat# A29127) were used for expression of recombinant proteins as described above.

SARS-CoV-2 (GISAID accession EPI_ISL_406862) was kindly provided by Bundeswehr Institute of Microbiology, Munich, Germany. Stocks were prepared by infection of Vero E6 cells for two days. When viral cytopathic effect (CPE) was observed, media were collected, clarified by centrifugation, aliquoted and stored at −80°C. Titer of stock was determined by plaque assay using Vero E6 cells.

SARS-CoV-2, isolate Human 2019-nCoV ex China strain BavPat1/2020 that was kindly provided by Prof. Dr. Christian Drosten (Charité, Berlin, Germany) through the European Virus Archive – Global (EVAg Ref-SKU: 026V-03883). The original viral isolate was amplified by 5 passages and quantified by plaque titration assay in Vero E6 cells, and stored at −80°C until use. The viral stock DNA sequence and coding capacity were confirmed as recently reported^59^.

Handling and working with SARS-CoV-2 was conducted in BL3 facility in accordance with the biosafety guidelines of the IIBR.

### Plaque reduction neutralization test (PRNT)

For plaque reduction neutralization test (PRNT)^60^, Vero E6 cells were seeded overnight (as detailed above) at a density of 0.5e6 cells/well in 12-well plates. Antibody samples were 2-fold serially diluted (ranging from 100 to 0.78 μg/ml) in 400 μl of MEM supplemented with 2% FBS, MEM non-essential amino acids, 2 nM L-glutamine, 100 Units/ml penicilin, 0.1 mg/ml streptomycin and 12.5 Units/ml Nystatin (Biological Industries, Israel). In cases where the PRNT applied for sera evaluation, 3-fold serially dilutions (ranging from 1:50 to 1:12,150) were used. 400 μl containing 300 PFU/ml of SARS-CoV-2 virus (GISAID accession EPI_ISL_406862) were then added to the mAb solution supplemented with 0.25% guinea pig complement sera (Sigma, Israel) and the mixture incubated at 37°C, 5% CO_2_ for 1 h. Monolayers were then washed once with DMEM w/o FBS and 200 μl of each mAb-virus mixture was added in duplicates to the cells for 1 h. Virus mixture w/o mAb was used as control. 2 ml overlay [MEM containing 2% FBS and 0.4% tragacanth (Sigma, Israel)] were added to each well and plates were further incubated at 37°C, 5% CO_2_ for 48 h. The number of plaques in each well was determined following media aspiration, cells fixation and staining with 1 ml of crystal violet (Biological Industries, Israel). Half-maximum inhibitory concentration (IC_50_) was defined as mAb concentration at which the plaque number was reduced by 50%, compared to plaque number of the control (in the absence of Ab).

### Animal experiments

Treatment of animals was in accordance with regulations outlined in the U.S. Department of Agriculture (USDA) Animal Welfare Act and the conditions specified in the Guide for Care and Use of Laboratory Animals (National Institute of Health, 2011). Animal studies were approved by the local ethical committee on animal experiments (protocol number M-57-20). Female K18-hACE2 transgenic (B6.Cg-Tg (K18-hACE2)2Prlmn/J HEMI; Jackson Laboratories, USA) were maintained at 20−22 °C and a relative humidity of 50 ± 10% on a 12 hours light/dark cycle, fed with commercial rodent chow (Koffolk Inc.) and provided with tap water *ad libitum*. All animal experiments involving SARS-CoV-2 were conducted in a BSL3 facility.

Infection experiments were carried out using SARS-CoV-2 BavPat1/2020 strain (EVAg Ref-SKU: 026V-03883). SARS-CoV-2 virus diluted in PBS supplemented with 2% FBS (Biological Industries, Israel) was used to infect animals by intranasal instillation of anesthetized mice. For mAbs protection evaluation, mice were treated by single administration of 1 ml volume, containing 0.1 and 0.01 mg Ab/mouse (equivalent of 5, and 0.5 mg/kg body weight, respectively) of the antibody 2 days following infection with 300 PFU SARS-CoV-2. Control groups were administered with PBS. Body weight was monitored daily throughout the follow-up period post infection. Mice were evaluated once a day for clinical signs of disease and dehydration. Euthanasia was applied when the animals exhibited irreversible disease symptoms (rigidity, lack of any visible reaction to contact).

## Results

### Phage-display library construction and screening for isolation of anti-NTD antibodies

Blood samples obtained from seven severe COVID-19 patients, were quantitatively evaluated by determining the antibody titers against S1 (by ELISA) and viral neutralization activities (by plaque reduction neutralization test, PRNT). The presence of anti-S1 antibodies (titers >10,000), as well as high neutralization potency (titers>1,000), were confirmed in all serum samples. In light of their high neutralization titers and in order to obtain a diverse antibody library, all seven samples were combined for the purpose of antibody library construction.

A phage-display single chain Fv (scFv) library, consisting of 1.3×10^9^ distinct clones was constructed for the subsequent antibody isolation. Three enrichment steps of panning were carried out by allowing the binding of phages to monomeric NTD, followed by screening of individual clones. ELISA of 400 individual clones from the 3^rd^ cycle, for NTD or RBD binding, resulted in the identification of >300 (75%) clones specific for NTD binding. One hundred and forty (140) clones were further examined by restriction fragment size polymorphism (RFLP), allowing identification of 19 unique patterns indicative of diverse sequences. Following DNA sequencing of clones representing these diverse patterns, 12 scFvs, showing sequence diversity, were expressed as scFv-Fc antibodies for further characterization. Sequence analysis was performed to the 12 mAbs, using IgBlast^25^, and germline families were assigned for each of the mAbs (Table 1).

**Table 1:** Germline usage and mutation frequencies of the isolated NTD-specific mAbs.

| mAb | IGHV | IGHD | IGHJ | IGLV | IGLJ | CDR3 length<br>(H,L) | SHM*<br>(H,L) |
| --- | --- | --- | --- | --- | --- | --- | --- |
| BLN1 | IGHV1-24 | IGHD1-20 | IGHJ6 | IGKV2-24 | IGKJ2 | 21,9 | 6,11 |
| BLN2 | IGHV1-24 | IGHD3-10 | IGHJ6 | IGLV2-14 | IGLJ2 | 21,10 | 3,10 |
| BLN3 | IGHV1-24 | IGHD1-20 | IGHJ6 | IGLV2-14 | IGLJ2 | 21,12 | 11,12 |
| BLN4 | IGHV4-30 | IGHD3-22 | IGHJ3 | IGLV1-44 | IGLJ1 | 21,11 | 5,6 |
| BLN5 | IGHV1-24 | IGHD1-20 | IGHJ6 | IGLV2-23 | IGLJ3 | 21,11 | 12,14 |
| BLN6 | IGHV4-61 | IGHD3-22 | IGHJ3 | IGLV2-8 | IGLJ1 | 21,10 | 4,12 |
| BLN7 | IGHV1-24 | IGHD6-19 | IGHJ5 | IGLV2-14 | IGLJ1 | 14,10 | 4,9 |
| BLN8 | IGHV3-30 | IGHD6-19 | IGHJ5 | IGLV3-1 | IGLJ2 | 10,9 | 4,3 |
| BLN9 | IGHV1-24 | IGHD6-19 | IGHJ5 | IGLV2-14 | IGLJ2 | 14,12 | 5,13 |
| BLN10 | IGHV1-24 | IGHD6-13 | IGHJ5 | IGLV3-19 | IGLJ2 | 16,10 | 4,22 |
| BLN12 | IGHV1-24 | IGHD1-20 | IGHJ6 | IGKV2-24 | IGKJ4 | 21,9 | 8,3 |
| BLN14 | IGHV3-30 | IGHD3-3 | IGHJ6 | IGKV3-20 | IGKJ5 | 16,8 | 4,1 |

DNA sequences of the antibodies were analyzed by IgBlast, for determining the germline allele usage and CDR3 loop length<sup>25</sup>.
\* No. of somatic hypermutations (SHM) relative to the respective V germline gene

The most prevalent V gene distinguished in the sequence of the mAbs was IGHV1-24, previously reported to be significantly enriched in anti-SARS-CoV-2 antibodies^23^. This germline family was also shown to be significantly enriched in expanded IgG+ B cell clones of COVID-19 patients, suggesting preferential recruitment and viral epitope binding by B cells exhibiting antigen receptors involving this IGHV allele^26^. Interestingly, BLN14 exhibited sequences originating from germline families IGHV3-30, IGKV3-20 and IGHJ6, which were previously shown to be over-represented in anti-SARS-CoV-2 mAbs^15^. This is especially notable considering that the combination of VH and VL did not originate in the context of the humoral immune response of the exposed individuals, but was rather the result of combinatorial pairing of heavy and light chains during library construction. This observation implies that combination of these particular germline families which was favorably selected in the course of screening, may preferentially generate antibodies exhibiting high affinity against SARS-CoV-2, as previously suggested^15,27^.

All 12 mAbs exhibited a relatively long CDRH3 loop (an average of 18 amino acids compared to an average of 13 for human antibodies^28^), a characteristic which was observed for many anti-SARS-CoV-2 mAbs, and in particular of non-RBD neutralizing antibodies^15,23^,26. In addition, the mAbs selected here reveal low levels of IGHV somatic hypermutations (SHM), 8 out of 12 exhibiting 5 or less mutations, and all twelve inspected exhibiting an average of 5.8 ± 2.9. This phenomenon was also observed in many SARS-CoV-2 mAbs and IgG+ B cell clones derived from COVID-19 patients^23,26^. This low level of SHM was shown to increase over time only at a moderate frequency^27^, indicating that the mutations level is independent of the time of isolation after the onset of the disease. The number of SHM was higher and more diverse in the IGLV genes (average of 9.7 ± 5.8), suggesting that this portion of the antibodies is subjected to a reduced level of selection constrains.

### Anti-NTD antibodies specificity, affinity and *in vitro* neutralization potency

Antibodies specificity to the NTD was confirmed by ELISA against NTD, RBD, S1 or spike proteins. All mAbs exhibited high specificity for NTD, and no binding to RBD. As expected, they also recognized both S1 and spike proteins (Fig. 1a).

**Fig. 1:**
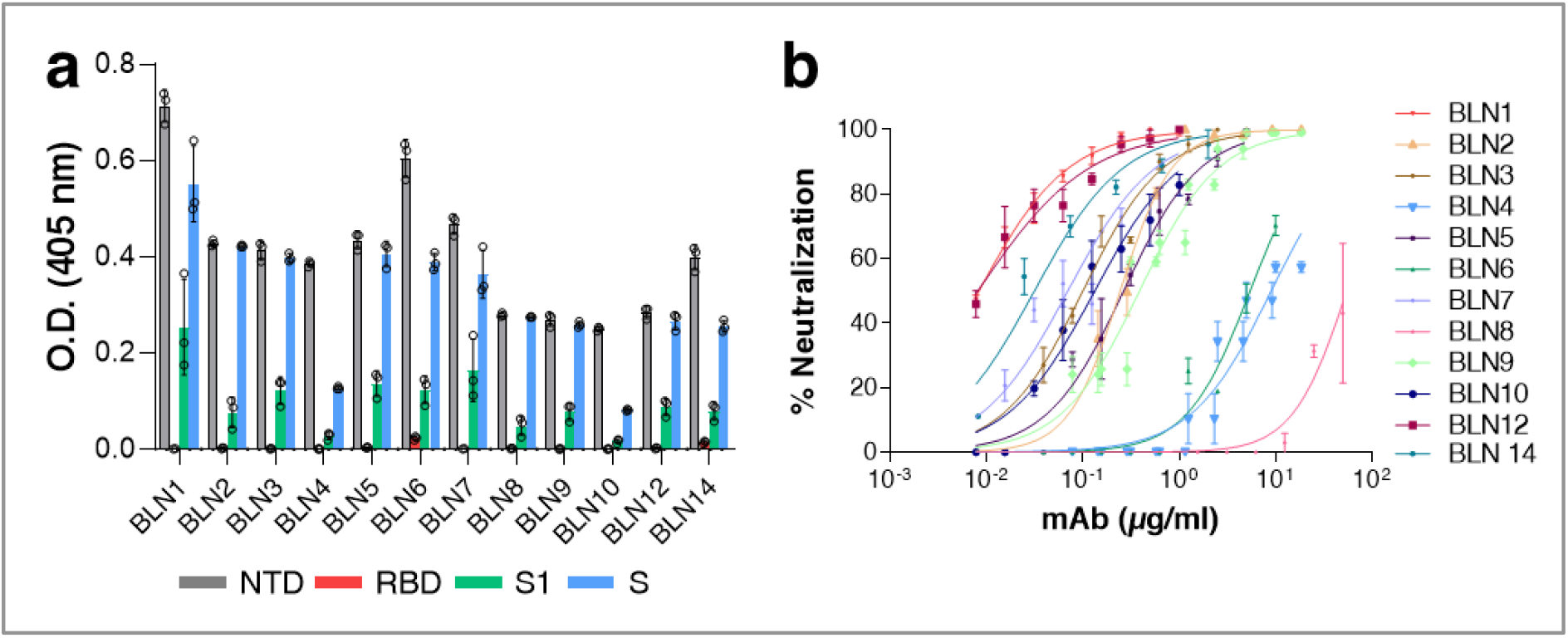
Characterization of individual mAbs isolated by phage display. **a-b**.Binding specificity and neutralization profiles of 12 selected anti-NTD mAbs (referred to as BLN1-BLN14). **a**. The specificity of each mAb was tested by ELISA using either SARS-CoV-2 S, NTD, RBD or S1 purified recombinant proteins as capturing antigens, as indicated by different colors. Data represent individuals and average of triplicates ±SEM. **b**. SARS-CoV-2 *in vitro* neutralization potency of the selected anti-NTD mAbs was evaluated by PRNT. Values along the curve depict averages of triplicates ± SEM. Each antibody is depicted by a different color, as indicated.

The affinity of each mAb for NTD was next determined, by Biolayer Interferometry (BLI). The K_D_ values calculated for each antibody, are presented in Table 2. Affinities ranged from 0.7 nM to 34 nM, BLN1 and BLN12 showing the highest affinity and BLN4, the lowest.

**Table 2:** Binding kinetic parameters of the selected NTD-specific mAbs, determined by BLI.

| mAb | K<sub>on</sub> (1/Ms) | K<sub>off</sub> (1/s) | K<sub>D</sub> (mM) |
| --- | --- | --- | --- |
| BLN1 | 2.47E+05 | 1.57E-04 | 0.7 |
| BLN2 | 1.81E+05 | 2.53E-03 | 14 |
| BLN3 | 3.14E+05 | 5.75E-04 | 1.8 |
| BLN4 | 1.19E+05 | 4.12E-03 | 34 |
| BLN5 | 2.63E+05 | 3.72E-04 | 1.4 |
| BLN6 | 1.94E+05 | 3.07E-03 | 15.8 |
| BLN7 | 2.86E+05 | 7.77E-04 | 2.7 |
| BLN8 | 1.35E+05 | 5.83E-04 | 4.3 |
| BLN9 | 2.35E+05 | 2.81E-03 | 12 |
| BLN10 | 2.11E+05 | 9.08E-04 | 4.3 |
| BLN12 | 3.63E+05 | 3.31E-04 | 0.9 |
| BLN14 | 9.31E+04 | 1.18E-03 | 12.7 |

The neutralization potency of the antibodies was evaluated by plaque reduction neutralization test (PRNT) using VeroE6 cells infected with authentic SARS-CoV-2. Accordingly, a fixed amount of the virus was incubated with increasing concentrations of each antibody, the mixture was then added to the cells, and the half-maximum inhibitory concentration (IC_50_) was determined 48 h later (Fig. 1b). All 12 antibodies were found to efficiently neutralize the virus, exhibiting IC_50_ values ranging from 54.9 µg/ml for BLN8, to a highly potent values of 0.008 µg/ml for mAbs BLN1 and BLN12 (supplementary table 1).

### Epitope mapping and determining the involvement of S1-appended *N*-glycans

The panel of antibodies was classified into clusters, representing competitive epitope recognition, as determined by BLI epitope binning assays (Fig. 2). Each individual antibody was biotinylated, immobilized to a streptavidin sensor, loaded with S1 and then competed with each of the other antibodies. The ability of two antibodies to bind S1 concomitantly (as identified by a wavelength shift in the interference pattern), indicates that they bind non-overlapping distinct epitopes^29^. Of note, the inability of two antibodies to bind together (competing antibodies) does not necessarily indicate that they bind the same epitope, as their competition may suggest steric interference due to binding to adjacent epitopes. Thus, the assay allows classification of antibody binding sites on the S1 subunit, based on the proximity of their respective epitopes. A representative example, showing sensograms of the 12 mAbs interactions with the pre-complex BLN12-S1, is depicted in Fig. 2a. As shown, antibodies BLN4, BLN6 and BLN8 recognize epitopes distinct from those recognized by BLN12, as manifested by a marked wavelength shift. However, all other antibodies did not elicit any marked shift in the presence of BLN12, indicating that they target overlapping or close epitopes. The ability of each antibody pair to simultaneously bind S1 was next determined for all the 11 other antibodies (Fig. 2b-binning matrix; supplementary Fig. 2). In summary, the 12 anti-NTD antibodies could be classified into three groups, based on recognition of three distinct epitopes. Group I includes BLN8, Group II includes BLN4 and BLN6 and Group III includes all other antibodies (Fig. 2c).

**Fig. 2:**
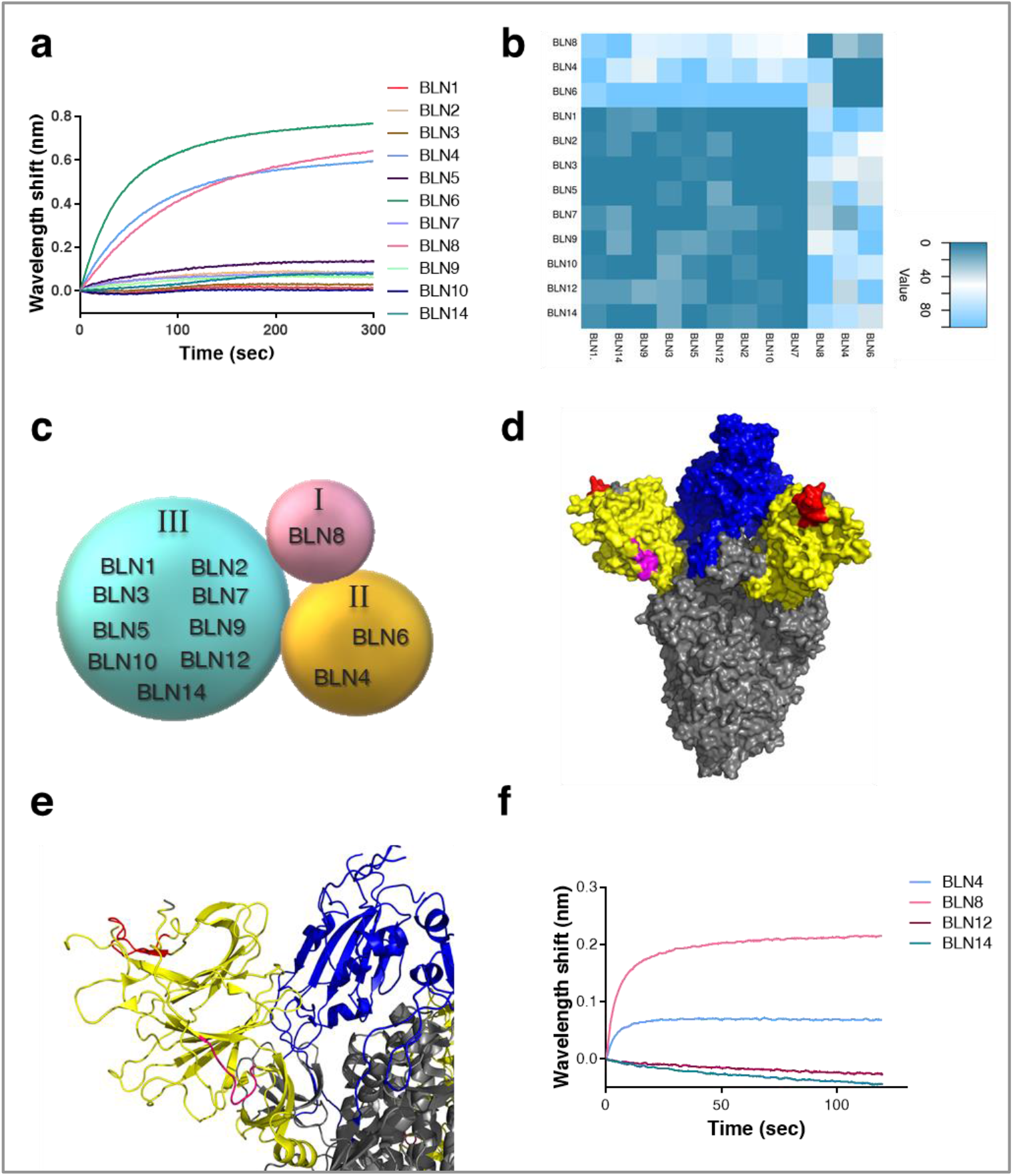
Epitope mapping of the selected 12 NTD-specific mAbs, determined by Biolayer interferometry (BLI). Each purified antibody was biotinylated, immobilized on a streptavidin sensor and loaded with S1. The complex (mAb-S1) was then incubated for 300 s with each one of the other antibodies. Distinct epitope recognition was evaluated by the ability of each pair of antibodies to simultaneously bind S1. **a**. Representative BLI results are depicted for the BLN12 mAb. Time 0 represents the binding to the BLN12-S1 complex by each of the indicated antibodies. Competition with antibody BLN12 itself served as a negative control, subtracted from the data before generation of the depicted sensograms. Detailed competition profiles obtained with all 12 antibodies are presented in Supplementary Fig. 2. **b**. Complete epitope binning results presented as a heat-map, indicating the normalized percentage values of area under curve (AUC) for each competition combination as calculated by Prizma (version 8.1). Normalization was performed individually for each capture antibody, 100% indicating lack of competition. Extent of competition between the capture antibody and the other antibodies, is represented by the indicated color-scale. Three distinct epitopes could be defined. **c**. Schematic interpretation of results from **a**, representing the classification of the selected mAbs into 3 groups (I-III). **d-e**. Space filling surface (**d**) and Ribbon (**e**) conformational models showing the location of the peptides identified by epitope mapping as the cognate epitopes of BLN4 and BLN12, on the surface of the viral spike protein [Protein Data Bank (PDB) ID 7C2L^24^]. RBD is depicted in blue, NTD in yellow and other regions in grey; BLN4-recognized peptide is depicted in pink and that of BLN12 in red. **f**. Competitive binding of representative mAbs with antibody 4A8, by BLI. Time 0 represents the binding of 4A8 to the indicated mAb-S1 complex.

Of note, the antibodies in group III (9 out of 12 mAbs), exhibited the highest neutralization potency compared to mAbs belonging to groups I and II, suggesting that the domain recognized by these antibodies, represents a major site of vulnerability of the SARS-CoV-2 and therefor a potential target for its neutralization.

In an attempt to further characterize the mAb specific recognition sites, representative members of the three groups were selected and subjected to epitope mapping, employing an array of overlapping synthetic peptides derived from the entire S1 protein. This strategy inherently enables only identification of linear epitopes, recognized by an antibody, therefore recognition of conformational epitopes which is often exhibited by neutralizing antibodies, cannot be detected. Indeed, mAbs BLN14 and BLN8 did not bind any of the peptides in the array. On the other hand, mAbs BLN4 and BLN12 were found to bind specific peptides, resulting in the identification of their recognition epitopes. The peptide recognized by BLN4 spans amino acids 56-65 (LPFFSNVTWF) of the spike protein (according to GenPept QHD43416 ORF) and the peptide recognized by BLN12 spans amino acids 141-155 (LGVYYHKNNKSWMES). The position of both mAbs on the spike structure is visualized in Fig. 2d-e. Notably, the identified peptide, found to be recognized by mAb BLN12, resides within the NTD N3 loop that was recently reported to be recognized by mAb 4A8, as determined by resolved structure of the 4A8 in complex with the SARS-CoV-2 spike^24^. In order to confirm the location of BLN12 epitope, a recombinant in-house version of 4A8 was generated and examined in an epitope binning assay together with the selected set of mAbs (Fig. 2f). Each of the selected mAbs was immobilized on a streptavidin sensor, loaded with NTD and further challenged with 4A8. Antibody 4A8 was indeed found to compete with mAbs BLN12 and BLN14, belonging to Group III. This competition further confirms BLN12 recognition site and reinforce the previous observation indicating an adjacent epitope for BLN14 as well. Interestingly, both BLN4 and BLN12 recognition sites, include *N*-glycosylation sites, located at residues N61 and N149 of the S protein, respectively^30^. Recognition of the synthetic peptides may imply that glycosylation (or other post-translational modifications) are not involved in the binding of these epitopes by the antibodies, yet the presence of the glycosylation signals encouraged inspection of glycans impact on the mAb interaction with the NTD. SARS-CoV-2 spike protein have been shown to express diverse *N*-glycans, including oligo-mannose and sialic acid-containing hybrid/complex-*N*-glycans^30^. Therefore, the four antibodies, representing the three groups, were examined on an exhaustive nano-printed glycan microarray, in which individual glycans are covalently immobilized via a primary amine linker onto an epoxide-coated glass slide. The glycan library contained a wide collection of glycans with terminal sialic acid (Sia), non-sialylated glycans (non-Sia)^31^, and a collection of homo- and hetero-glycodendrons of mono/oligo-mannose and -galactose ligands (Man)^32^ (Supplementary Fig. 3). While there was very low binding to most examined glycans, the antibodies revealed a strong preference to the *N*-glycans-associated structures LNT-3Ac (Glycan ID #60; Neu5Acα3Galβ3GlcNAcβ3Galβ4GlcβOR) and LNT (Glycan ID #59; Galβ3GlcNAcβ3Galβ4GlcβOR) (Supplementary Fig. 3). However, binding was not detected when the terminal sialic acid was replaced with Neu5Gc (*N*-glycolylneuraminic acid), the hydroxylated form of Neu5Ac (LNT-3Gc; Glycan ID #61; Neu5Gcα3Galβ3GlcNAcβ3Galβ4GlcβO-R) (Supplementary Fig. 3). To further investigate these interactions, the antibodies were subjected to a competition assay assessing inhibition of antibodies binding to NTD, using glucuronic acid (GlcA), Neu5Ac in its closed ring α-linked structure (2-*O*-methyl-α-Neu5Ac; Ac2Me), or with sialoglycopeptides^33^ containing a mixture of *N*/*O*-glycans with a terminal Neu5Ac^34^. Both GlcA and Ac2Me showed no inhibition of antibodies binding, however the Neu5Ac-sialoglycopeptides completely abrogated binding of BLN4 and BLN12, but not that of BLN8 and BLN14 (Fig. 3a), suggesting that the interactions with the *N*-glycans are involved in the binding to NTD of BLN4 and BLN12 in particular.

**Fig. 3:**
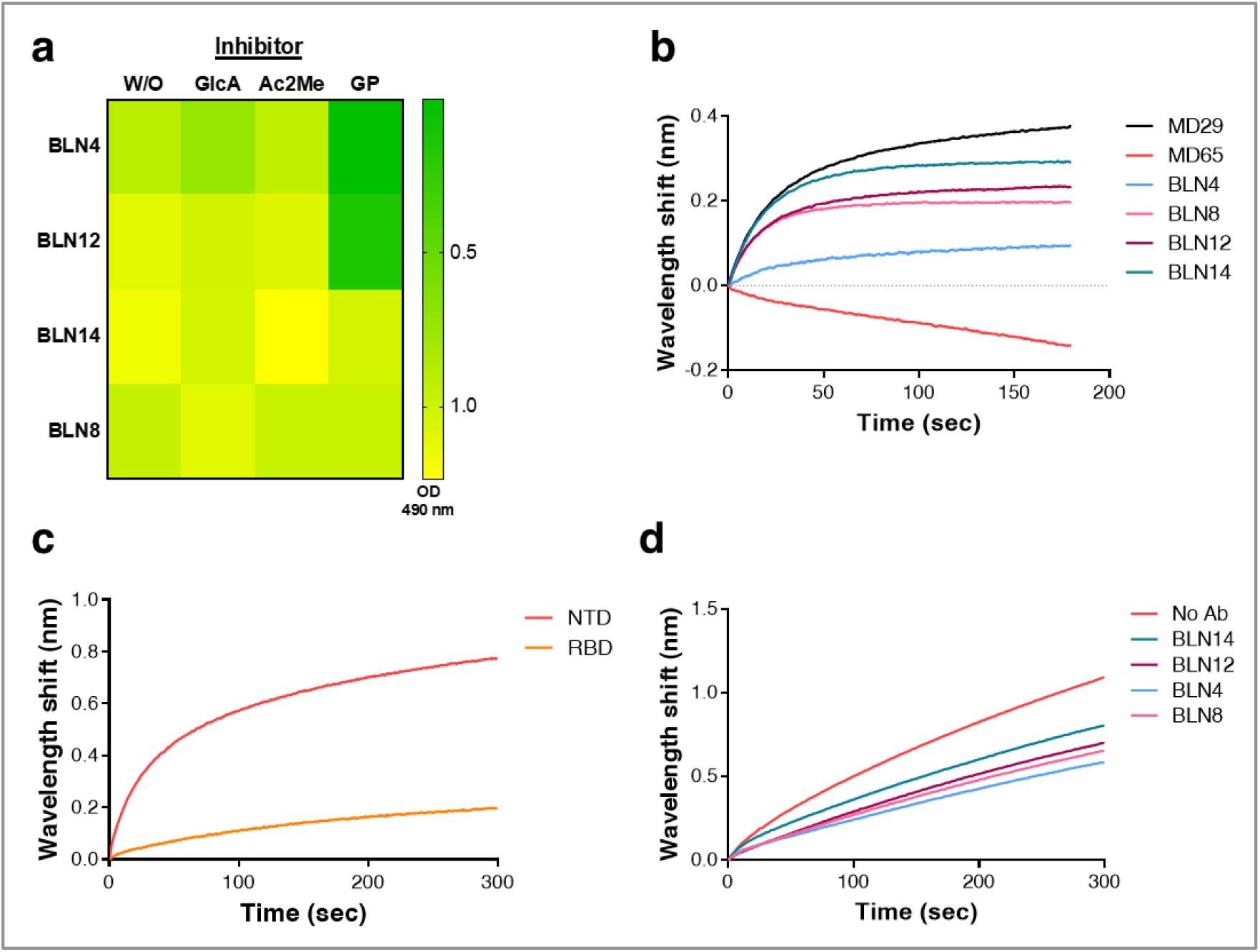
Evaluation of the involvement of hACE2, L-SIGN and various glycan structures in the neutralization promoted by mAbs. **a**. Competition of mAbs binding to NTD by various glycans was determined by ELISA. Antibodies were pre-incubated on ice in the presence or absence of: 2-*O*-methyl-α- Neu5Ac (Ac2Me; 8 mM), D-glucuronic acid (GlcA; 8 mM), or with sialoglycopeptides containing a mixture of *N*/*O*-glycans with a terminal Neu5Ac (GP; 0.06 mM). Inhibition is manifested by prevention of binding, resulting in a low level of yellow coloring. None of the antibodies were inhibited by GlcA or Ac2Me, while BLN4 and BLN12 were inhibited by the GP. **b-d**. The ability of purified recombinant hACE2 and human L-SIGN receptors to bind S1 or NTD, in the presence of antibodies was tested by BLI. **b**. Binding of hACE2 to S1 in the presence of neutralizing antibodies (representing each of the epitope groups, depicted by different colors, as indicated). Each of the biotinylated antibodies was immobilized on a streptavidin sensor, loaded with S1, washed and incubated with recombinant hACE2 for 300 s. Time 0 represents the binding of the hACE2 to the antibody-S1 complex. **c**. Binding of human L-SIGN to NTD and RBD. L-SIGN-Fc was immobilized on a protein A sensor and incubated with NTD or RBD. **d**. Binding of human L-SIGN to NTD in the presence of neutralizing antibodies (representing each of the epitope groups). NTD-His was immobilized on Ni-NTA sensors and saturated with each of the antibodies separately; the NTD-mAb complex was then incubated with L-SIGN for 300 s. Time 0 represents the binding of the L-SIGN to the antibody-NTD complex.

These results are in-line with the epitope mapping data and collectively suggest that BLN4 and BLN12 recognize both the amino-acid sequences and the glycan moieties appended to their cognate respective epitopes on NTD.

### Addressing the neutralization mechanism of selected anti-NTD mAbs

To address the mechanism underlining the neutralization characteristics of the anti-NTD mAbs, a hACE2 binding inhibition assay was conducted, as blockage of the hACE2 receptor engagement is considered to represent the most effective virus neutralization mechanism of anti-RBD mAbs. Thus, biotinylated-BLN antibodies were immobilized on streptavidin sensors and their ability to inhibit S1 binding to hACE2 was tested by BLI (Fig. 3b). A wavelength shift indicates binding of hACE2 to mAb-S1 complex and hence, lack of inhibition by the mAb tested. Four mAbs, representing the three epitope groups were examined. The anti-RBD antibodies MD65, previously demonstrated to inhibit hACE2 binding, and MD29 which does not inhibit hACE2 binding^19^, were used as positive and negative controls, respectively.

Results indicate that, as anticipated, none of the tested anti-NTD mAbs inhibits hACE2 binding. It may therefore be confirmed that the neutralization mechanism promoted by these antibodies does not involve the inhibition of hACE2 binding.

It was recently reported that in addition to the “canonical” ACE2 receptor binding by RBD, SARS-CoV-2 NTD interacts with C-type lectin receptors, including L-SIGN and DC-SIGN^11,13^ as an alternative route for SARS-CoV-2 infection and entry into host cells. Accordingly, the ability of four representative anti-NTD mAbs (which bind different epitopes on the NTD) to inhibit the interaction of the NTD with L-SIGN, was examined (Fig. 3c-d). Binding of NTD to L-SIGN was first evaluated in comparison to RBD using BLI. Indeed the NTD was found to interact with L-SIGN, while RBD showed only low level of interaction, as also reported previously^11,12^ (Fig. 3c). This interaction, was then evaluated in the present of selected anti-NTD mAbs (Fig. 3d), showing partial inhibition, ranging from approximately 25% binding inhibition by BLN14 to 50% inhibition by BLN4. We may therefore conclude that neutralization promoted by the anti-NTD antibodies partially involves interference with the binding of L-SIGN receptor and that additional entities or functions may be involved.

### Evaluating the therapeutic potential of anti-NTD mAbs in a SARS-CoV-2 lethal infection model

For *in vivo* assessment of the therapeutic potential of the antibodies, the anti-NTD BLN12 and BLN14 were selected. Antibody BLN12 was superior to the other antibodies in the panel, in terms of affinity (0.9 nM) and *in vitro* neutralization potency (IC_50_=8 ng/ml). Although antibody BLN14 exhibited lower affinity (12.7 nM) than others, it revealed relatively high *in vitro* neutralization (IC_50_=30 ng/ml) and its sequence differed from that of BLN12 (Supplementary Table 1; Table 1). Antibodies BLN12 and BLN14 were therefore examined for their ability to protect K18-hACE2 transgenic mice from an authentic SARS-CoV-2 infection *in vivo*. The K18-hACE2 transgenic murine model was shown to be highly susceptible to SARS-CoV-2 infection, resulting in significant viral load in the lungs, heart, brain and spleen, as well as high mortality^35–37^. This model was recently shown to fatefully recapitulate the SARS-CoV-2 infection and consequently to serve as a reliable model to predict the efficacy of therapeutic strategies^35,38 36,38,39^. In a recent study, we applied this model for demonstration of the protection efficacy of therapeutic mAb directed to the S1-RBD, when administered 2 days post-infection (dpi)^14^.

For the *in vivo* experiments, BLN12 and BLN14 were produced as full recombinant IgG molecule of the IgG1/k isotype which includes the triple-mutation M252Y/S254T/T256E (YTE) in the Fc region. These modifications are intended to prolong the antibodies serum half-life in humans (by increasing their affinity towards the human FcRn), a parameter that is essential for high therapeutic value^40–42^.

K18-hACE2 transgenic mice were infected with 300 pfu/mouse SARS-CoV-2, resulting in more than 75% mortality within 7-10 days following infection. Based on the superior *in vitro* neutralization ability of the BLN12 and BLN14 anti-NTD mAbs, the *in vivo* potency of these two mAbs was directly tested at a stringent model of 2 dpi. (Fig 4). Each antibody was administered as a single dose of either 0.1 or 0.01 mg/mouse infected with 300 pfu SARS-CoV-2. Body weight and survival of experimental animals were monitored daily for 21 days post-infection. While a mortality rate of 75% was observed in control mice (infected but untreated with antibodies), administration of 0.1 mg/mouse of BLN14, induced full protection and prevented any weight loss (Fig. 4a-b). Even at 0.01 mg dose, BLN14 treatment resulted in survival of ~85% of the animals and the survivors recovered and regained their initial weight by day 10 post-infection.

**Fig. 4:**
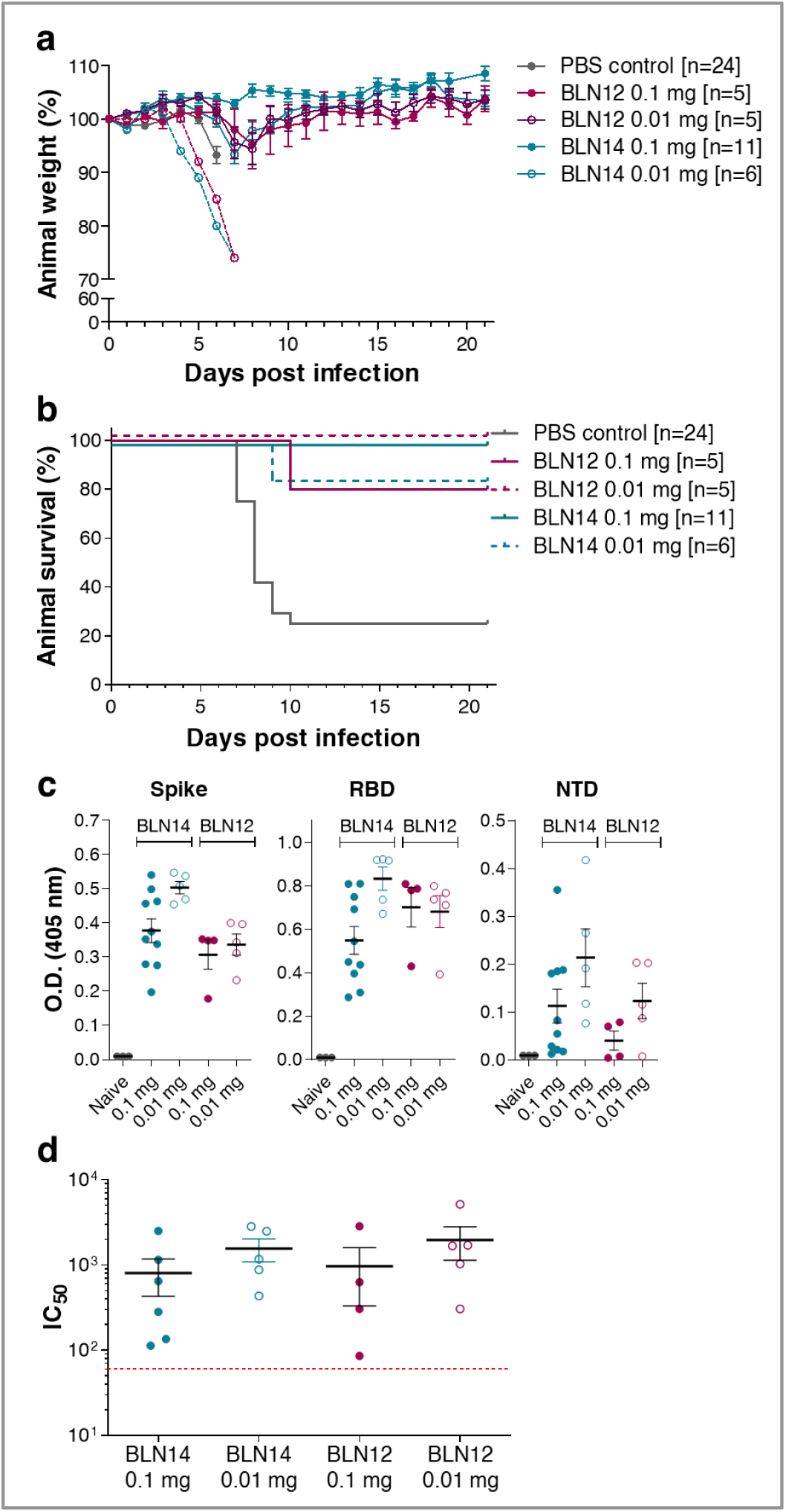
Post-exposure therapeutic efficacy of BLN12 and BLN14 in SARS-CoV-2 infected K18-hACE2 mice. Single doses of 0.1 or 0.01 mg Ab/animal, were administered at day 2 post viral infection. **a**. Body weight profiles. Body weight is displayed as percentage of initial weight (colored differently according to the mAb and the dose used, as indicated within the panel). Only data of the first 7 days is presented in the control group exhibiting significant mortality. The weight of the animals which died in the experimental groups (BLN14 0.01 mg [n=1] and BLN12 0.1 mg [n=1]) is indicated by separate dashed line. **b**. Kaplan-Meyer surviving curves. The mAb and the dose used, are indicated within the panel. **c**. Endogenous anti-SARS-CoV-2 humoral response of infected K18-hACE2 mice treated with the BLN12 or BLN14 antibodies. Sera samples were collected at day 21 post-infection from mice treated with BLN12 or BLN14 at 2 dpi with the indicated doses, and tested by ELISA for the presence of endogenous (murine) antibodies recognizing the SARS-CoV-2 antigens: spike (left panel), RBD (central panel) or NTD (right panel). Data represent individual and mean ±SEM of measurements obtained for sera samples at 1:100 dilutions. **d**. SARS-CoV-2 *in vitro* neutralization potency of the sera samples detailed in **c**, evaluated by PRNT. Data represent individual and mean ±SEM of IC_50_ Values. Dotted red line indicates the assay limit of detection.

Treatment with BLN12 resulted in a similar protection rate, consisting of 80% and 100% survival after treatment with 0.1 mg and 0.01 mg, respectively. Mice experienced only a minor weight loss and recovered by day 10 post-infection. These results indicate a high efficacy of both mAbs *in vivo*, even at low doses and at a late time point post infection.

### Endogenous humoral response to SARS CoV-2 of antibody-treated animals

Administration of antibodies enables efficient therapy both by neutralization of the pathogen, as well as by extending the time window necessary for the mounting of an endogenous host response. Therefore, elicitation of an endogenous humoral response following passive immunization, represents a significant advantage of mAb treatment, especially in case where re-infection is a tangible possibility, as in the case of the SARS-CoV-2 pandemic. Hence, the murine humoral response against SARS-CoV-2 related antigens was measured in BLN12- and BLN14-treated mice (Fig. 4c). Serum samples were collected 21 dpi following post-exposure (2 dpi) administration of each mAb, at dose of 0.1 or 0.01 mg per animal. The murine antibody response was tested by ELISA against SARS-CoV-2 spike glycoprotein, RBD and NTD (Fig 4c). Note that in this timepoint, no excess of the human mAb, used for treatment, could be detected. All treated mice revealed elicitation of immune responses against the three tested antigens. This result is in a good agreement with our previous observation, using anti-RBD mAb^14^. In this previous report, endogenous response, following treatment with high mAb dose, was shown to correlate with time of treatment, demonstrating a marked response in post-exposure treatment, rather than in prophylactic one. In the current study, by evaluating the 2 doses of mAb treatment at 2 dpi, a significant endogenous response is demonstrated (Fig 4c). Furthermore, this antibody response was demonstrated to consist of significant neutralizing activity, as was demonstrated by applying PRNT (Fig. 4d).

## Discussion

The continuing escalation of the COVID-19 pandemic, complicated by the recent emergence of SARS-CoV-2 variants, motivate the development of efficient means for therapeutic intervention. Here we described the isolation of a panel of novel recombinant human mAbs, targeting the SARS-CoV-2 NTD domain originating from libraries constructed with antibodies collected from COVID-19 active patients. The germline usage, epitope specificities, binding characteristics and neutralizing abilities of 12 antibodies were determined; subsequent assessment of two of them *in vivo* revealed exceptional potency promoting full protection of lethally infected mice and suggesting that they may serve as promising candidates for antibody-mediated therapy. In combination with anti-RBD mAbs, therapeutic administration of these antibodies may circumvent complications related to the occurrence of viral escape mutants which may not be neutralized by targeting only one particular epitope.

Germline usage analysis of the 12 mAbs isolated in the current study, determined high preference for germline genes also observed for other SARS-Cov-2 antibodies^15,23^. Dominant germlines in antibodies targeting RBD were previously reported^43^ therefore the present data imply that similar dominant germlines are used for generation of anti-NTD antibodies as well. The anti-NTD mAbs exhibited low numbers of somatic mutations, similarly to anti-RBD mAbs, and the frequency of somatic mutations could not be correlated to neutralization potency or to a certain epitope, suggesting that they did not occur as a result of selection for high antigen affinity. Interestingly, anti-RBD mAbs were shown to possess a relatively short CDRH3, while most anti-NTD mAbs display a significantly longer CDRH3, a difference which probably reflects the dissimilar nature of the epitopes: while anti-RBD antibodies recognize surface epitopes, the NTD epitopes reside within a large pocket in the spatial architecture of the S1 molecule, to which antibodies exhibiting longer CDRH3 loops may preferentially bind efficiently. Of note, BLN8, representing a unique epitope amongst the novel mAbs, was the only one displaying a relatively short CDRH3. These observations are in line with those recently reported by Yuan et al.^43^ showing that a short CDRH3 represents a characteristic molecular features common to the antibody response to SARS-CoV-2 RBD, possibly owing to the fact that the specific epitopes recognized by these antibodies are relatively flat.

Many viruses, including HIV-1^44,45^ and SARS-CoV-2^30^, display a strong glycan shield as they bud out of the infected host cells, possibly as a host cell mimicking mechanism, for evading the immune system. It has been shown that some of the most potent neutralizing antibodies against HIV-1 target this glycan shield^45^, not by recognizing a shallow groove within a binding pocket, but rather by a mechanism allowing for such antibodies to be “embraced” by extended glycan chains. In particular, the PG9 antibody is uniquely characterized by an unusually long CDRH3 loop of its heavy chain forming a hammerhead-like structure accommodated by two *N*-glycan arms extending out from the viral gp120 trimer (within the highly variable V1/V2 domain). This phenomenon contributes to the stabilization of the interaction between the gp120 trimer and the antibody facilitating neutralization of the virus^46,47^. Here we show that some of the SARS-CoV-2 neutralizing antibodies characterized by a long CDR3 (i.e. BLN4, BLN12), similarly interact with extended *N*-glycan structures of NTD.

One of the important aspects of the antibodies presented here is that they neutralize the SARS-CoV-2 virus by a mechanism which does not involve interference with engagement of the ACE2 receptor, therefore their potency underlines the importance of alternative modality employed by the virus primary interaction with the target host cell at the onset of infection. Although SARS-CoV-2 mainly infects the lungs, it also targets multiple other organs including the cardiovascular system, gastrointestinal tract, and kidneys^48–50^, tissues where hACE2 expression is very low or undetectable, implying such an alternative infection route. The C-type lectin family is one of the largest subgroup of the lectin superfamily. These proteins possess carbohydrate-recognition domains, recognizing specific carbohydrate structures and thus mediate cell-cell and cell-pathogen interactions^51^. Such receptors are expressed at high level in cells of the human immune system, including monocytes, dendritic cells and macrophages, which were implicated in the pathogenesis of SARS-CoV-2^52,53^. The densely glycosylated SARS-CoV-2 spike protein was recently shown to directly interact with a variety of lectin receptors, such as DC-SIGN and L-SIGN, suggesting an alternative route for SARS-CoV-2 viral entry^11–13^.

Epitopes recognized by BLN12 and BLN14 were mapped to the same cluster, by the binning experiments, yet it was clearly demonstrated that they do not share exactly the same epitope. We show that while BLN12 identifies glycans as part of its epitope, BLN14 does not. Both antibodies also differ in their ability to bind linear epitopes and slightly differed in their capability to inhibit NTD binding to L-SIGN. Nevertheless, as explained, the epitopes recognized by these two antibodies are expected to be proximal. The two mAbs exhibited similar *in vitro* and *in vivo* neutralization capabilities. It is conceivable therefore to assume that their neutralization mechanism is very similar. Glycans on N149 of NTD were suggested to mainly mediate the interaction with L-SIGN, while *N*-glycosylation in positions N74 and N282 may have a partial role in such interaction^11^. Epitope mapping data pointed to N149 as a part of BLN12 epitope, and glycan inhibition assay confirmed the presence of glycans within this mAb epitope recognition site. Nevertheless, it could not completely inhibit L-SIGN binding to NTD. It can therefore be concluded that the neutralization of SARS-CoV-2 by BLN12 and BLN14 only partially rely on inhibition of virus binding to lectin receptors such as L-SIGN. This interference may be significant in the context of virus attachment to cells. The possibility that these mAbs prevent other interactions, such as membrane fusion or proteolytic activation cannot be ruled out.

Stabilization of one conformational state of viral fusion proteins, was suggested as a possible neutralization mechanism of mAbs, for different viruses such as MERS-CoV and HIV^54,55^. Since BLN12 and BLN14 showed very high *in vitro* and *in vivo* neutralization ability, without inhibiting binding to either hACE2 or L-SIGN receptors, this stabilization of an unfavorable conformation, provides a possible explanation for their mechanism of neutralization. Indeed, the inability of the spike protein to bind C-type lectins in the presence of antibodies, as was shown by cell assays^11^, may be the consequence of a conformational state imposed by the antibodies, rather than a direct interference to binding. If L-SIGN and DC-SIGN are indeed alternative receptors for SARS-CoV-2 and mediate its infection of cells in many tissues lacking hACE2, it may be speculated that anti-NTD mAbs may also exert their therapeutic activity by limiting the spreading of the virus in the body. This aspect further emphasizes the importance of including anti-NTD mAbs in antibody-based therapeutics for SARS-CoV-2, in addition to its contribution in circumventing the complication of escape mutant occurrence. The observation that the anti-NTD mAbs provide full protection of K18-hACE2, a murine model of COVID-19 which relies on transgenic expression of the hACE2, may contradict the conclusion that the antibodies do not directly interfere with the binding of the virus to hACE2 (as emerging from the binding assays). This discrepancy may be resolved by assuming that the interaction of the anti-NTD antibodies with the spike protein occurs prior to the engagement of the virus to the hACE2. Such an interaction may neutralize the virus by a variety of manners such as expediting its clearance and/or interfering with some adherence mechanism of the virus to the target cell, necessary for facilitating the engagement to the hACE2 receptor. In any event, full understanding of the phase of infection affected by the therapeutic antibodies and the co-activity of alternative internalization mechanisms of the virus into the host cells will require further studies.

In conclusion, the 12 mAbs reported here, recognized three distinct epitopes within the spike-NTD. Two of these epitopes, represented by BLN4 and BLN8, seem to be less important for the virus neutralization. The third epitope, however, was found to be a major point of vulnerability for the virus. Antibodies BLN12 and BLN14, directed to this epitope, and characterized in this study, showed remarkable neutralization of the authentic SARS-CoV-2 *in vitro* and more importantly, allowed full protection in an animal model of lethal infection, even when administered at a low dose (0.5 mg/Kg) and at a late time point after infection (2 dpi). Notably, although the high susceptibility of the K18 transgenic mouse model to SARS-CoV-2 infection is associated with the hACE2 expression, treatment with NTD specific mAbs (that are not interrupting with the hACE2 interaction), resulted in equivalent protection efficacy, observed by the potent RBD specific mAb MD65, as reported lately^14^. This observation further emphasize the complementary protection efficacy of NTD-specific mAbs. The BLN12 and BLN14 mAbs, reported herein therefore, represent excellent candidates for therapy of SARS-CoV-2, possibly in combination with anti-RBD mAbs.

## Acknowledgments

We thank Prof. Dr. Christian Drosten at the Charité Universitätsmedizin, Institute of Virology, Berlin, Germany for providing the SARS-CoV-2 BavPat1/2020 strain. We wish to express our gratitude to our colleagues Dr. Tomer Israely, Dr. Nir Paran, Dr. Sharon Melamed, Boaz Politi, Dr. Emanuelle Mamroud, Dr. Hadar Marcus, Dr. Noa Madar-Balakirski, for fruitful discussions and support. We would like to thank Moshe Mantzur, Tamar Aminov, Keren Eden and Noam Mazuz for skillful and devoting technical assistance. We thank Mr. Mark Power, from the British Embassy in Tel-Aviv for his highly appreciated assistance in facilitating the research collaboration involving COVID-19 patients’ blood samples. We would like to express our appreciation to Einat Ben-Arie, Sarit Ohayon, Yael Shlomo and Ronen Zolshain for intensive administrative support.

## Author Contributions

T.N-P., A.M., Y.L., E.M., R.A., D.G., M.A., R.F., S.L., A.Z., E.E, S.W., H.A., O.M. and R.R. designed, carried out and analyzed the data. S.L.B-A, performed the glycan analyses. J. D. E., R.K., H.Y. and X.C. provided crucial reagents. T.C., S.Y., S.S. and V.P-K added fruitful discussions, reviewed and edited the manuscript. R.R. and O.M. supervised the project. All authors have approved the final manuscript.

## Data and materials availability

Antibodies are available (by contacting Ohad Mazor from the Israel Institute for Biological Research) for research purposes only under an MTA, which allows the use of the antibodies for non-commercial purposes but not their disclosure to third parties. All other data are available from the corresponding author upon reasonable requests.

## Supplementary Information

### Highly potent therapeutic antibodies targeting the SARS-CoV-2 spike N-terminal domain

Tal Noy-Porat *et. al*

## Supplementary Table

**Supplementary Table 1.**
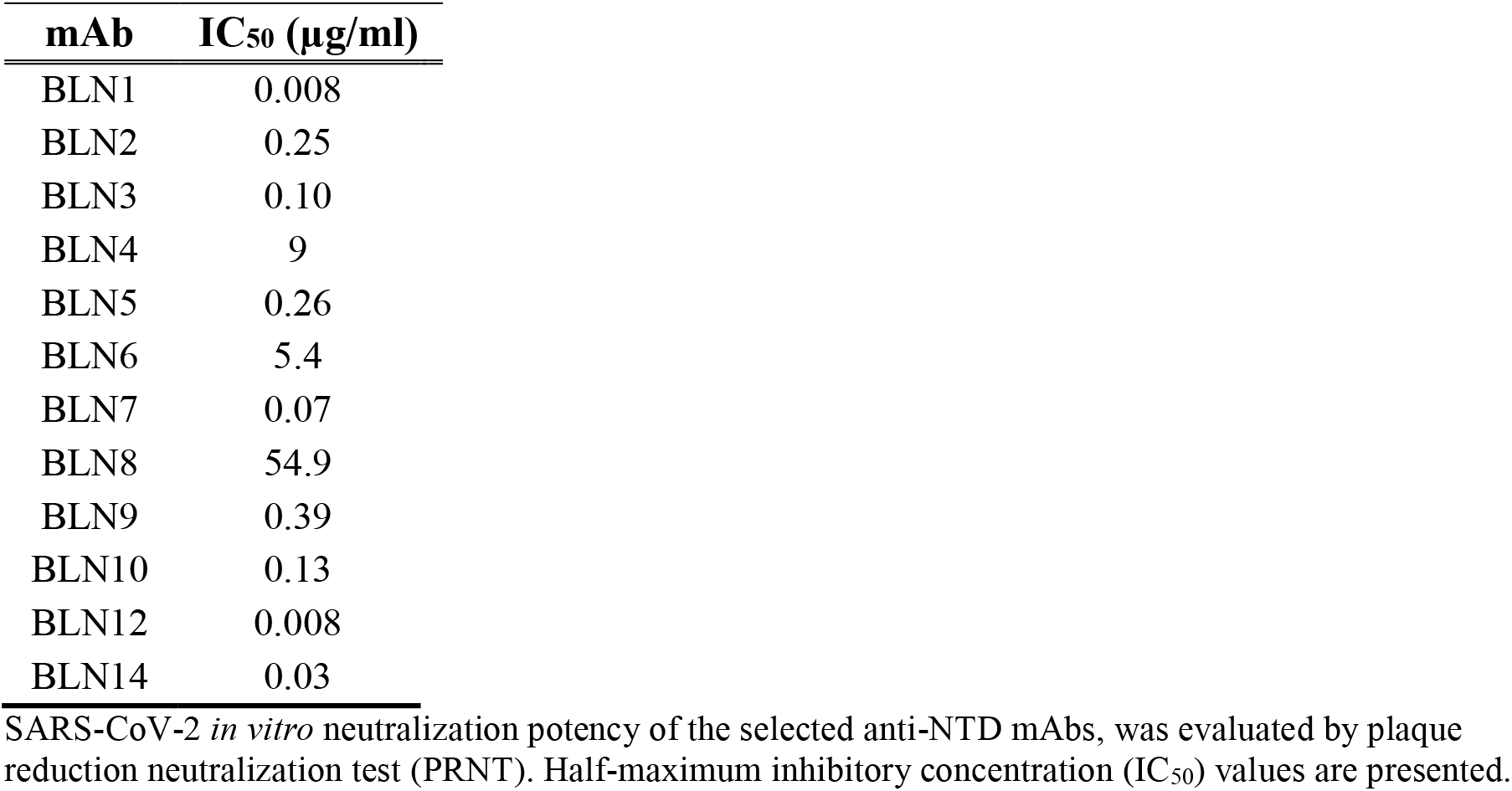
*In vitro* neutralization of SARS-CoV-2 by NTD-specific mAbs.

| **mAb** | **IC<sub>50</sub> (µg/ml)** |
| --- | --- |
| BLN1 | 0.008 |
| BLN2 | 0.25 |
| BLN3 | 0.10 |
| BLN4 | 9 |
| BLN5 | 0.26 |
| BLN6 | 5.4 |
| BLN7 | 0.07 |
| BLN8 | 54.9 |
| BLN9 | 0.39 |
| BLN10 | 0.13 |
| BLN12 | 0.008 |
| BLN14 | 0.03 |

SARS-CoV-2 *in vitro* neutralization potency of the selected anti-NTD mAbs, was evaluated by plaque reduction neutralization test (PRNT). Half-maximum inhibitory concentration (IC<sub>50</sub>) values are presented.

## Supplementary Figures

**Supplementary Figure 1.**
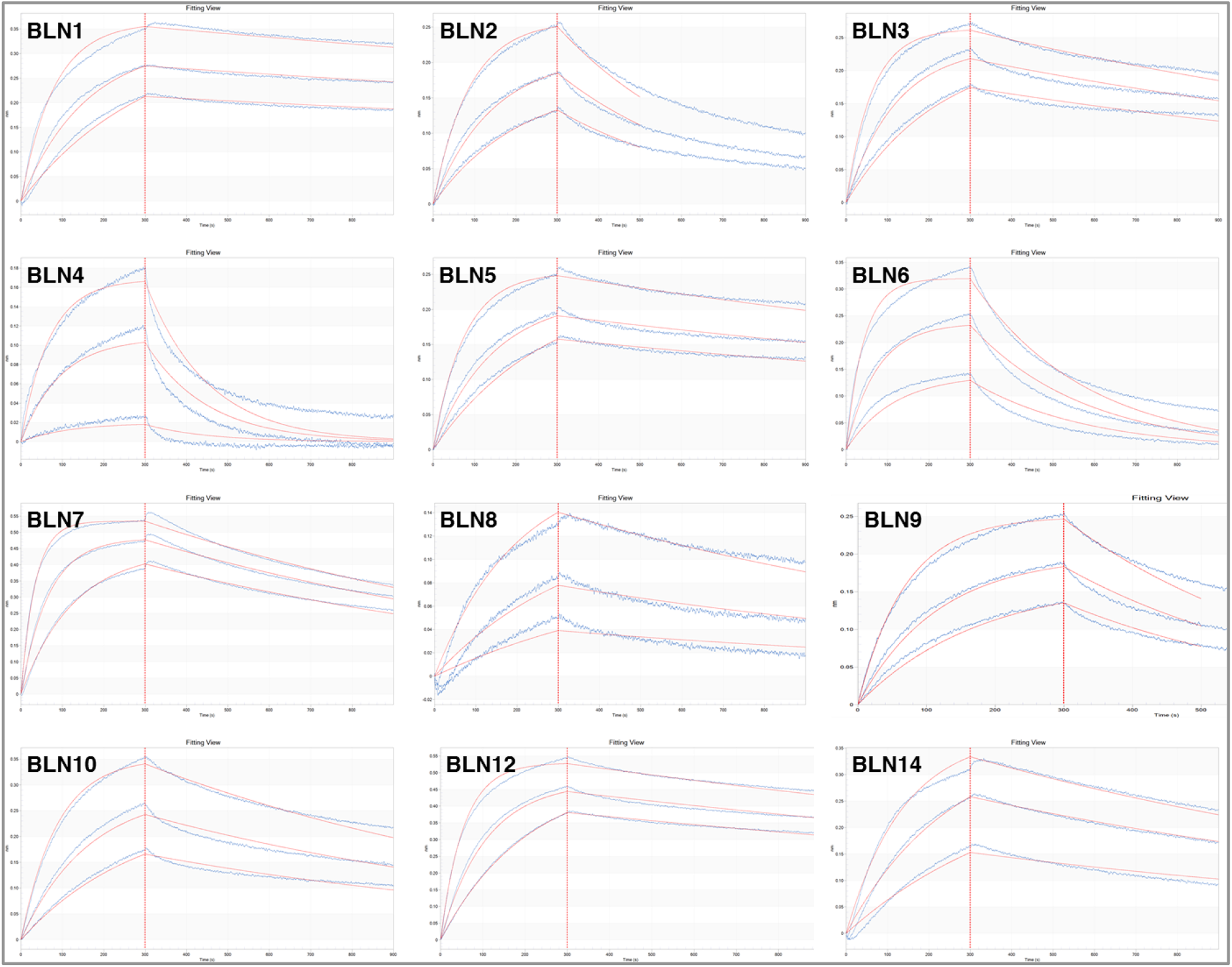
NTD-specific mAbs affinity determination by BLI. Biolayer interferometry analysis used for the determination of the 12 selected anti-NTD mAbs. Streptavidin-coated biosensors were loaded with each biotinylated mAb and reacted for 300 s with the indicated concentrations of the recombinant RBD (association phase) and then transferred to buffer-containing wells for another 600 s (dissociation phase). Sensograms (after subtraction of parallel measurements from unloaded biosensors) were fitted with 1:1 binding model (red curves) using the Octet data analysis software 8.1.

**Supplementary Figure 2.**
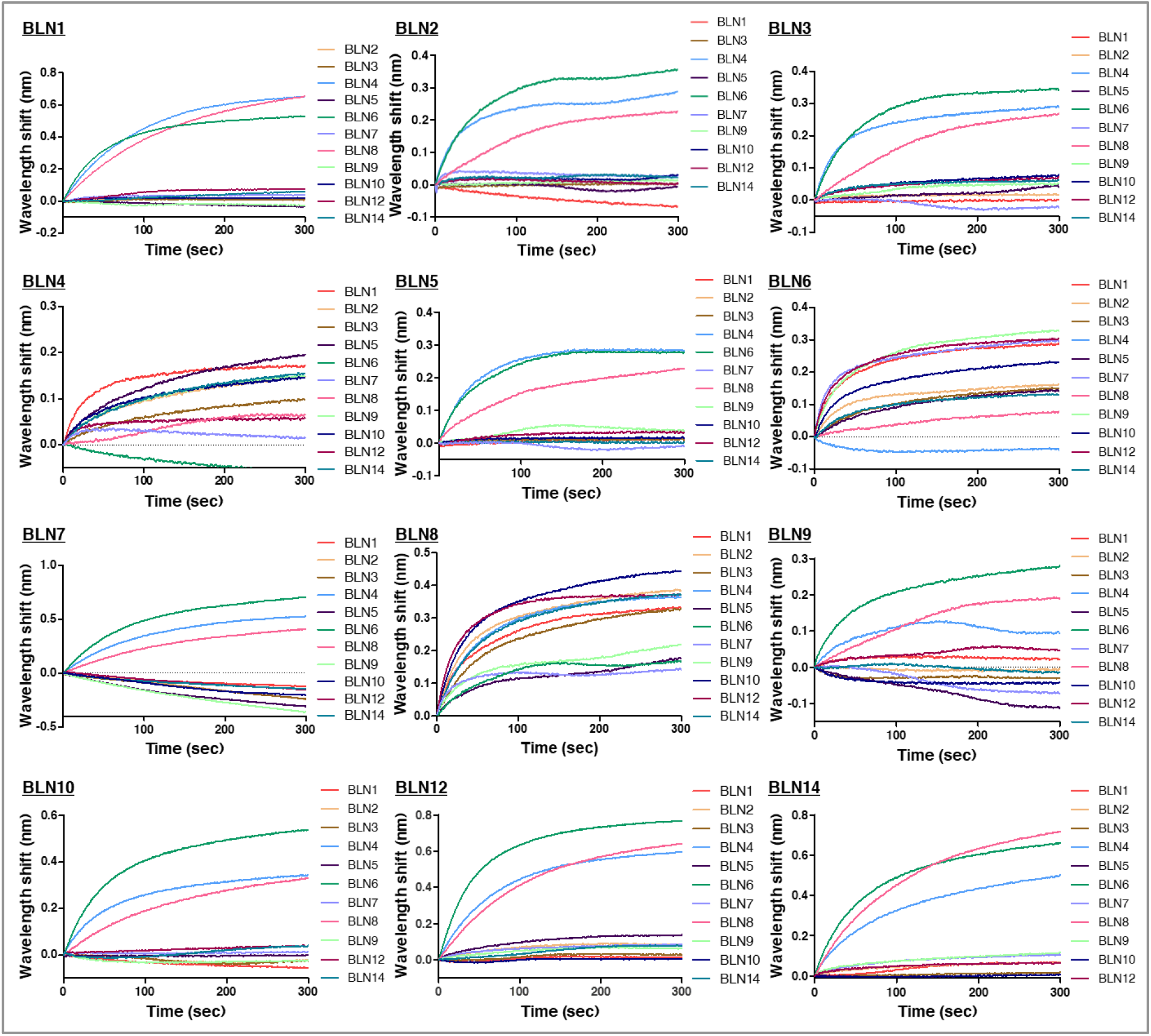
Epitope binning of the selected anti-NTD mAbs, by Biolayer interferometry (BLI). Each purified antibody (indicated on the left upper side of each sensogram) was biotinylated, immobilized on a streptavidin biosensor and saturated with S1. The complex (mAb-S1; Time 0) was then incubated for 300 s with each one of the other antibodies (colored differently according to the mAb, as indicated within the panels). Competition of each mAb with itself, served as a negative control, subtracted from the data before generation of the depicted sensograms.

**Supplementary Figure 3.**
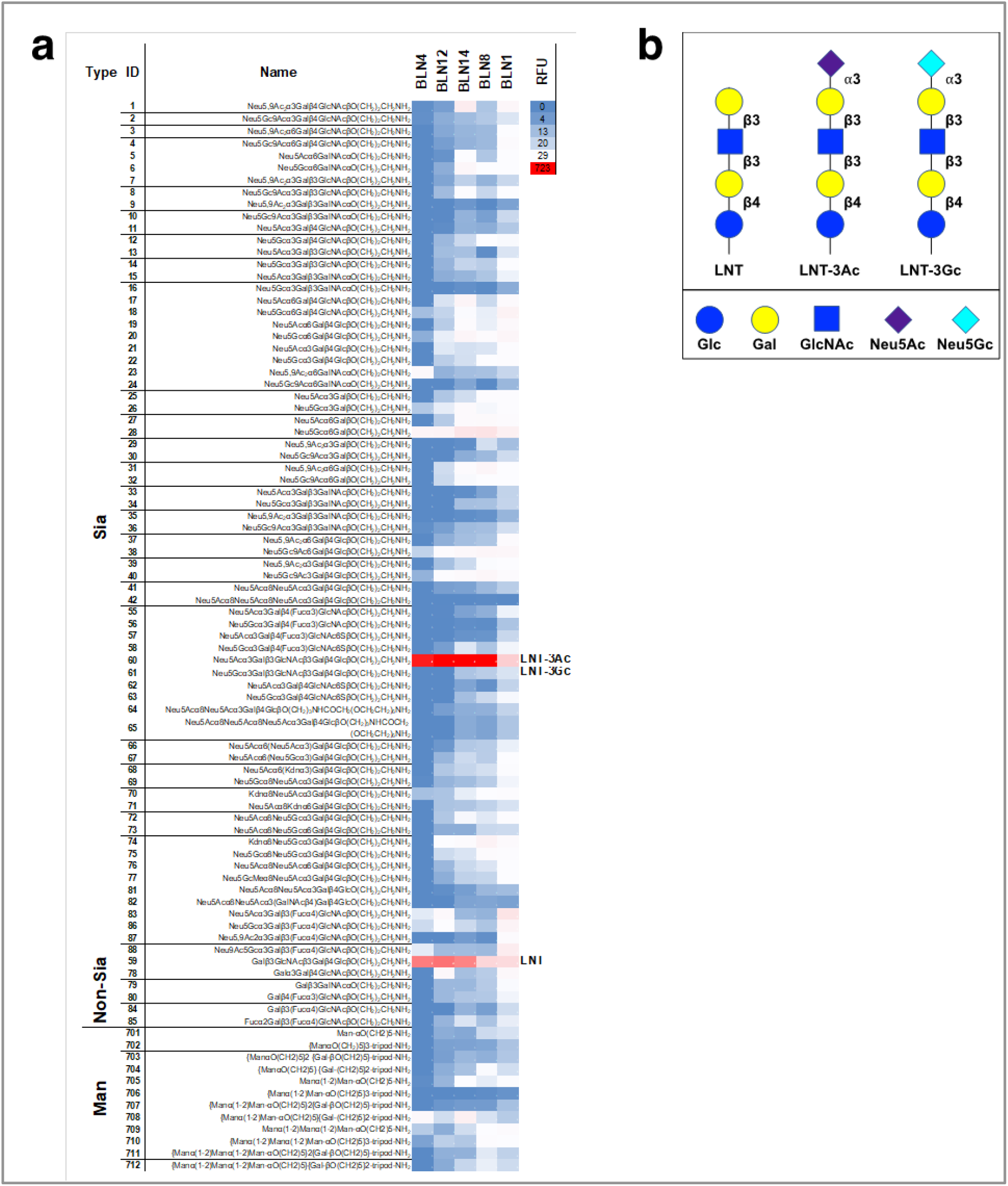
Antibodies glycan recognition. **a**. Antibodies glycan recognition was characterized by glycan microarrays against printed sialylated glycans (Sia), non-sialylated glycans (Non-Sia) or glycodendrons of mannose and galactose ligands (Man), detected with Cy3-goat-anti-human IgG. Arrays were scanned, analyzed using GenePix pro-7 and relative fluorescent units (RFU) were calculated. Binding is shown as a heatmap of all the arrays (red highest, blue lowest and white 80^th^ percentile). Binding was most prominent against *N*-glycan moieties LNT-3Ac and its non-sialylated derivative LNT, but not to LNT-3Gc. **b**. Schematic illustration of the glycans: LNT (Galβ3GlcNAcβ3Galβ4GlcβO-R), LNT-3Ac (Neu5Acα3Galβ3GlcNAcβ3Galβ4GlcβO-R) and LNT-3Gc (Neu5Gcα3Galβ3GlcNAcβ3Galβ4GlcβO-R).

